# The Length of Promoter Sequence Affects The *De Novo* Initiation By T7 RNA Polymerase in Vitro: New Insights Into The Evolution of Promoters For Single Subunit RNA Polymerases

**DOI:** 10.1101/619395

**Authors:** Ramesh Padmanabhan, Dennis Miller

## Abstract

RNA polymerases (RNAPs) differ from other polymerases in that they can bind promoter sequences and initiate *de novo* transcription. Promoter recognition requires the presence of specific DNA binding domains in the polymerase. The structure and mechanistic aspects of transcription by the bacteriophage T7 RNA polymerase (T7 RNAP) are well characterized. This single subunit RNAP belongs to the family of RNAPs which also includes the T3, SP6 and mitochondrial RNAPs. High specificity for its promoter, the requirement of no additional transcription factors, and high fidelity of initiation from a specific site in the promoter makes it the polymerase of choice to study the mechanistic aspects of transcription. The structure and function of the catalytic domains of this family of polymerases are highly conserved suggesting a common mechanism underlying transcription. Although the two groups of single subunit RNAPs, mitochondrial and bacteriophage, have remarkable structural conservation, they recognize quite dissimilar promoters. Specifically, the bacteriophage promoters recognize a 23 nucleotide promoter extending from −17 to + 6 nucleotides relative to the site of transcription initiation, while the well characterized promoter recognized by the yeast mitochondrial RNAP is nine nucleotides in length extending from −8 to +1 relative to the site of transcription initiation. Promoters recognized by the bacteriophage RNAPs are also well characterized with distinct functional domains involved in promoter recognition and transcription initiation. Thorough mutational studies have been conducted by altering individual base-pairs within these domains. Here we describe experiments to determine whether the prototype bacteriophage RNAP is able to recognize and initiate at truncated promoters similar to mitochondrial promoters. Using an *in vitro* oligonucleotide transcriptional system, we have assayed transcription initiation activity by T7 RNAP. When a complete or almost complete (20 to 16 nucleotide) double stranded T7 RNAP promoter sequence is present, small RNA’s are produced through template-independent and promoter-dependent stuttering corresponding to abortive initiation, and this effect was lost with a scrambled promoter sequence. When partial double stranded promoter sequences (10 to 12 nucleotides) are supplied, template dependent *de novo* initiation of RNA occurs at a site different from the canonical +1-initiation site. The site of transcription initiation is determined by a recessed 3’ end based paired to the template strand of DNA rather than relative to the partial promoter sequence. Understanding the mechanism underlying this observation helps us to understand the role of the elements in the T7 promoter, and provides insights into the promoter evolution of the single-subunit RNAPs.

## 1.2 Introduction

The evolution of genetic information depends in part on the coevolution of polymerases that can synthesize the informational molecule and at the same time transfer the genetic information in the template through the formation of Watson-Crick base pairing. The earliest polymerases in an RNA World are thought to be ribozymes with RNA-directed RNA synthesis. During the transition from the RNA-world to an RNA-protein world the ribozymes were replaced by single subunit protein polymerases with RNA-directed RNA synthesis [RNA replicase]. The transition from the RNA-Protein world through the RNA-DNA Retro world to the modern DNA world required a co-evolution of polymerase properties to give RNA-directed DNA synthesis [reverse transcriptases], and DNA-directed DNA polymerases [DNA polymerases, DNA Replicases], respectively. One argument that genetic information has evolved through these different “Worlds” is the modern existence of these coevolved single subunit polymerases. All of these single subunit polymerases presumably initiated synthesis through a primer extension mechanism, since almost all modern single subunit polymerases retain this property.

Polymerases can be classified into four categories based on the type of nucleic acids synthesized and templates used: 1] DNA-directed DNA polymerases (DNAPs; DdDp), 2] DNA-directed RNA polymerases (RNAPs; DdRp), 3] RNA-directed DNA polymerases (reverse transcriptases or RTs), and 4] RNA-directed RNA polymerases (RNA replicase; RdRp) (Ahlquist, 2002; Castro et al., 2009; Chen & Romesberg, 2014). These polymerases can be further divided into single subunit RNA polymerases and multi-subunit RNA polymerases (Werner & Grohmann, 2011). The single subunit RNAPs have motifs in common and are thought to derive from a common ancestor but the evolutionary divergence of these polymerases is still obscure (Cermakian et al., 1997).

A major evolutionary step was the evolution of the promoter which defined genes with specific functions from the genetic information, along with the coevolution of domains in RNA polymerases which would recognize promoter sequences. This innovation results in *de novo* initiation of transcription and provides a way to specifically regulate gene expression. RNAPs are distinguished by their ability to recognize promoter sequences and initiate transcription *de novo*, rather than extend from a primer (Cheetham & Steitz, 2000; Kennedy, Momand, & Yin, 2007; Steitz, Smerdon, Jager, & Joyce, 1994). This initiation phase is transient and generally occurs while the RNAP is bound to the promoter (Gong & Martin, 2006; Liu & Martin, 2002). The transition to the elongation phase requires a change in RNAP structure to allow promoter release and processive movement on the template (promoter clearance) (Gong, Esposito, & Martin, 2004; Martin, Muller, & Coleman, 1988). This initiation phase is unique to RNAPs and this ability to recognize promoters and initiate *de novo* is a key step in the evolution of organisms’ ability to transfer specific genetic information from DNA.

The bacteriophage T7 RNA Polymerase is the prototype single subunit RNA polymerase. It is an ideal model system for studying polymerase and promoter evolution. It is related to other bacteriophage RNA polymerases and the mitochondrial RNA polymerases. This group of RNAPs has very conserved structure and sequence. However, the mitochondrial RNA polymerases generally have different promoter sequences from the bacteriophage polymerases (Figure 2.1). Bacteriophage polymerases generally have a 23 nucleotide promoter that overlaps the site of transcription initiation by six nucleotides (−17 to + 6). Mitochondrial RNAPs recognize a diverse variety pf promoter sequences which are typically about nine nucleotides in length and run from - 8 to +1. In figure 1 we use the well characterized mitochondrial promoter from the yeast mtRNAP as a reference for comparison with the bacteriophage RNAPs. The yeast promoter sequence has similarity with the −8 to +1 portion of the T7 promoter.

**Figure 2.1.**
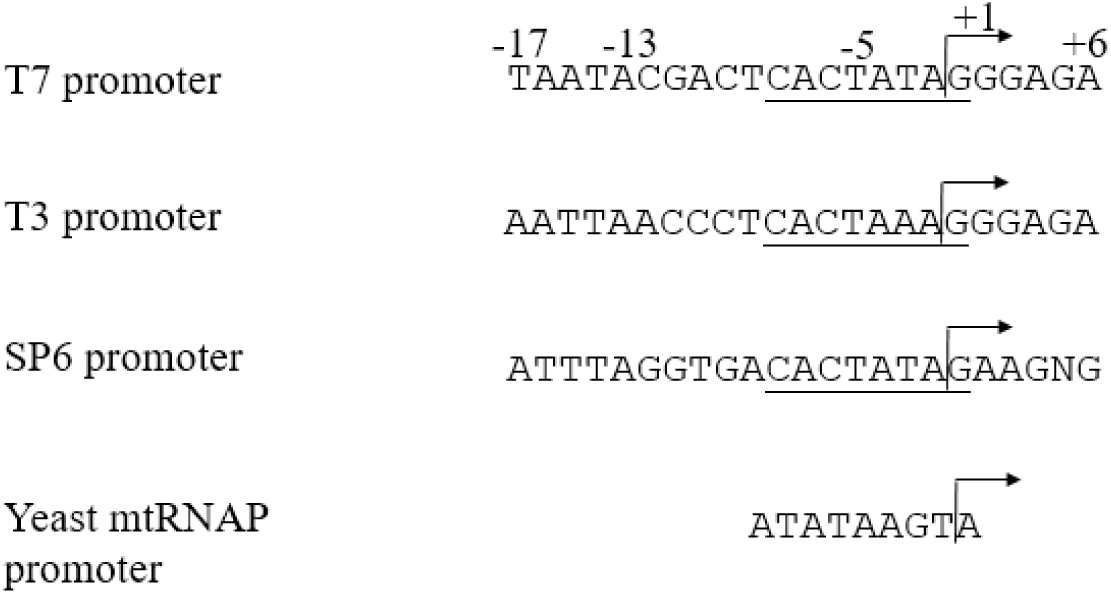
Comparison of three bacteriophage RNAP promoters and yeast mtRNAP (YmtRNAP) promoter. All sequences shown in the 5’-3’ orientation. The −8 to +1 portion of the bacteriophage promoter is underlined. The nine-nucleotide yeast mtRNAP promoter sequence is aligned with the bacteriophage −8 to +1 sequence to highlight their analogous positions relative to transcription initiation.

The 23 nucleotide promoter of the T7 RNA polymerase can be divided into several functional domains (Figure 2.2). The initiation region covers the ten nucleotides from - 4 to +6. Nucleotides +1 to +6 are called the transcription start site. The +1 site is conserved as a G in all the bacteriophage promoters. Sites +2 to + 6 are conserved as purines in the bacteriophage promoters. Polymerase contacts in this region are primarily with the pyrimidine, template strand. Substitution of these nucleotides decreases promoter strength and initiation efficiency only slightly. The region −4 to −1 is called the unwinding region. Positions −1 [A], −3 [A], −4 [T] are conserved among the bacteriophage promoters. This region is invariably AT-rich presumably to aid in melting of the DNA strands at the initiation site, although the conservation of positions −1, −3 and −4 may indicate polymerase contact sites.

**Figure 2.2.**
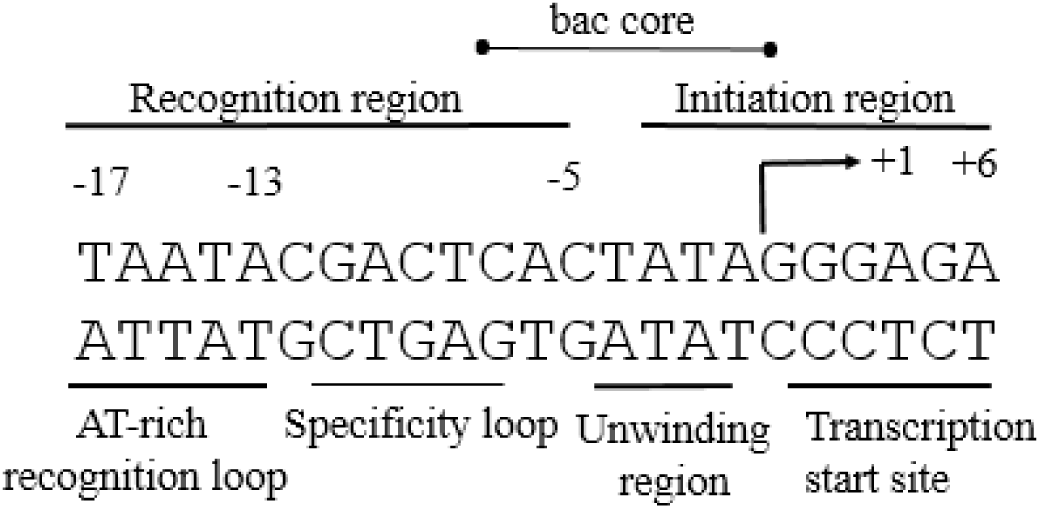
The T7 RNAP promoter showing functional domains

The promoter recognition region (13 nucleotides) extends from −5 to −17 and consist of polymerase interaction sites important for polymerase binding and positioning. These promoter contact sites interact with two regions of the polymerase. Promoter positions −5 to −12 interact with a T7 RNAP structural domain called the specificity loop, located near the carboxyl terminus, while positions - 13 to −17 interact with a T7 RNAP structural domain called the AT-rich recognition loop located near the amino terminus of the T7 RNA polymerase.

The phylogenetic tree depicting the evolutionary relationships among the single subunit RNAPs has not been rooted and so the evolutionary relationship between the two classes of promoter sequence is unknown (Cermakian et al., 1997). The T7 RNAP can recognize a range of promoter sequences which are closely related to each other by this consensus sequence (Dunn & Studier, 1983; Tang, Bandwar, & Patel, 2005). Details concerning the T7 RNAP promoter structure and function were deduced from studies on mutant promoter sequences (Ikeda, Ligman, & Warshamana, 1992; Klement et al., 1990). Single base changes either in the recognition region or the initiation domain affects the efficiency of promoter recognition and initiation of transcription respectively, but a mutation in the recognition region does not affect initiation and vice versa (Ikeda et al., 1992). The promoter region from −17 to −13 is dispensable, and can be deleted with no effect on initiation activity, however, optimal promoter recognition requires contacts within this region (Chapman & Wells, 1982; Martin & Coleman, 1987; Osterman & Coleman, 1981). Comparison of the promoter sequences from the three phage promoters T3, T7 and SP6 revealed that all the three promoters share a similar core sequence from −7 to +1 pointing to a common role of this region in the promoter function (Brown, Klement, & McAllister, 1986; Jorgensen, Durbin, Risman, & McAllister, 1991), Figure 2.1). There is considerable sequence divergence from −8 to - 12 corresponding to the specificity loop-recognition region in the T7 RNAP promoter, suggesting that this region plays a key role in sequence-specific contacts. Though there is an 82% homology in the amino acid sequence between T3 and T7 RNAPs, neither of the enzymes can efficiently initiate transcription at promoters used by the other (Klement et al., 1990; Rong, He, McAllister, & Durbin, 1998). Promoter specificity studies using base-pair substitutions at −10, −11, and −12 positions in the T7 RNAP promoter by residues of T3 RNAP promoter revealed that base pairs - 10 to −12 have important role with −11 base having a significant role in promoter binding (Klement et al., 1990). Base substitutions at positions −7 (A or G for C), −8 (A for T), −9 (A or T for C, −11 (T for G) completely inactivated the T7 promoter (Chapman, Gunderson, Anello, Wells, & Burgess, 1988). Methylation interference studies show that binding of T7 RNAP to the promoter was interfered by methylation of the G-residues at −7 and −9 in the template strand and −11 in the non-template strand suggesting that T7 RNAP makes important contacts in the major groove between −7 and −11 (Jorgensen et al., 1991).

The yeast *Saccharomyces cerevisiae* mitochondrial RNA polymerase (YmtRNAP) is homologous to the single subunit bacteriophage T3 and T7 RNAPs (Cermakian, Ikeda, Cedergren, & Gray, 1996; Masters, Stohl, & Clayton, 1987; Matsunaga & Jaehning, 2004). YmtRNAP recognizes the simple nine-nucleotide-long promoter consensus sequence ATATAAGTA for transcription initiation, which differs in sequence and length from the phage RNAPs (Nayak, Guo, & Sousa, 2009). However, this sequence can be aligned with the −8 to +1 core region of phage RNAPs (Figure 2.1). The YmtRNAP has a region analogous to the specificity region which by analogy interacts with −8 to −5 [ATAT]. However, there is no analogous region in yeast mtRNAP to the A-T rich binding region in T7RNAP and so the yeast promoter consensus does not extend beyond −8.

To study T7 RNAP’s ability to use truncated promoters similar to mitochondrial promoters, we developed an *in vitro* transcription system based on the *in vitro* transcription system developed by Milligan *et al.* (Milligan, Groebe, Witherell, & Uhlenbeck, 1987) and Weston et al. (Weston, Kuzmine, & Martin, 1997)They have shown that the 18 nucleotides, −17 to +1, when double stranded and attached to 5’ extended template is sufficient to act as a promoter in vitro (Figure 2.3A). When T7 RNAP is presented with these oligonucleotides and the four ribonucleotide triphosphates, they produce a run-off RNA starting with GGG. We have modified their procedure by extending the double stranded region to 20 nucleotides (−17 to +3) as in Figure 2.3B in order to increase initiation frequency. With T7RNAP and the four ribonucleotide triphosphates present, these oligonucleotides produce a run-off RNA of the same length starting with GGG. We further modified this model by creating an oligonucleotide which formed a intramolecular double stranded region of the type shown in Figure 2.3C. Under the same conditions it produced an RNA of the same length and composition.

**Figure 2.3.**
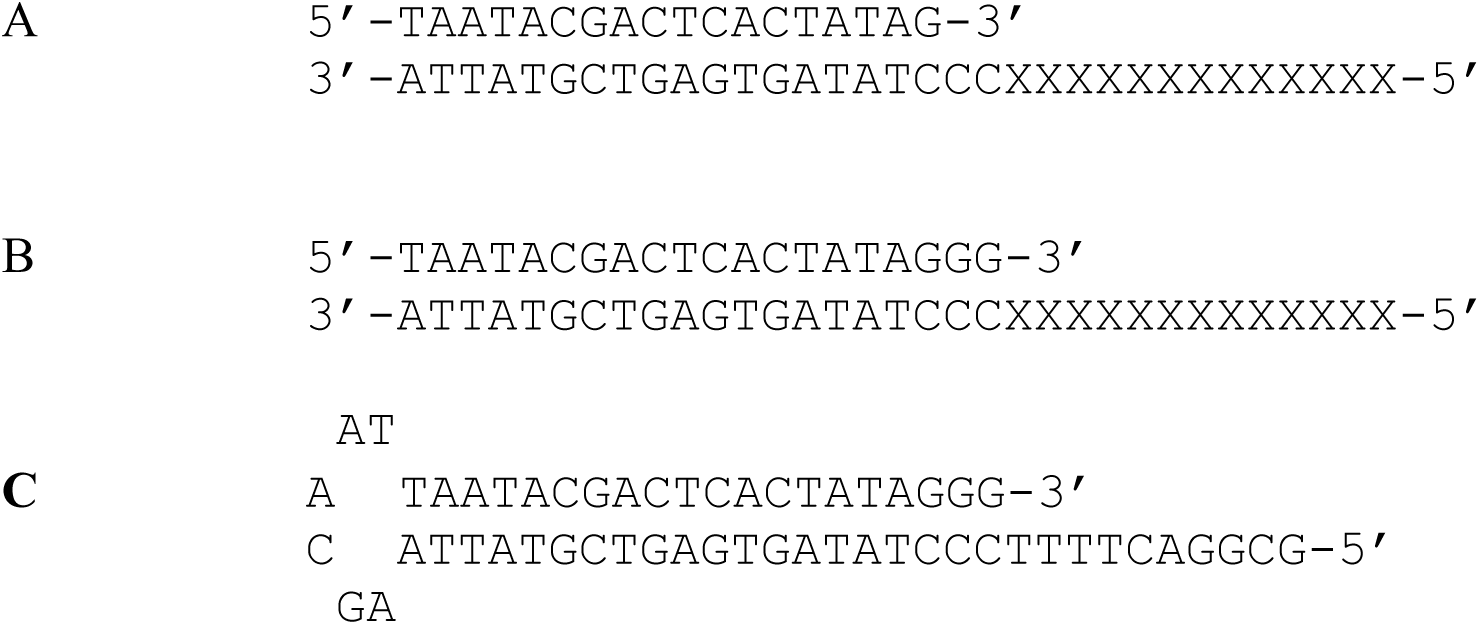
Oligonucleotides used for run-off transcription with T7RNAP. Panel A shows oligonucleotides used by (Milligan et al., 1987)) and (Weston et al., 1997) Panel B shows second type of oligonucleotide which can be used for runoff transcription. Panel C shows an oligonucleotide of the type used in this study to generate run-off RNAs. Xs indicate variable regions in which nucleotide substitutions can be made.

We have used this oligonucleotide system to determine whether T7 RNAP in the presence of such an oligonucleotide and a single ribonucleotide triphosphate can correctly and efficiently initiate transcription on truncated promoter sequences resembling mitochondrial promoters.Here, we show that T7 RNAP can initiate on truncated promoter sequences similar to mitochondrial promoters. However, the site of transcription initiation is not at the canonical initiation site, but initiates instead on the first unpaired base in the template. These results are analyzed in the light of T7 promoter flexibility, promoter evolution, and the use of this technique to produce defined RNAs without 5’- sequence constraints and 3’-variability.

## 1.3 Materials and methods

### 1.3.1 Oligonucleotides, Radiolabeled nucleotide triphosphate and T7 RNAP

Oligonucleotides designed for this study were procured from Eurofins MWG operon, USA. The desiccated oligonucleotides were resuspended in water to a concentration of 100μM. The radiolabeled α^32^P ATP was procured from Molecular Bioproducts, USA. T7 RNA polymerase enzyme (50,000 U/mL) was procured from New England Biolabs, USA.

### 1.3.2 Reaction conditions

50μL reaction mixtures containing 5μl of the 100 μM oligonucleotides, 2μL of radiolabeled α^32^P-ATP (3000Ci/mmol), 5μL of 10X RNA polymerase reaction buffer (1X concentrations of 40mM Tris-HCl, 6mM MgCl_2_, 2mM spermidine, 1mM dithiothreitol), supplied along with T7 RNA polymerase, 1μL of RNAase inhibitor-RNasin (Promega, USA) and 1μL (50U) of T7 RNA polymerase were incubated at 37°C for 60 minutes. Reactions were stopped and run on 15% Polyacrylamide gel at 75V. The gels were then stained with ethidium bromide to visualize the nucleic acids, and subsequently exposed to a phosphor screen (Amersham GE Healthcare) for 5 minutes and scanned using Storm 840 (Amersham GE Healthcare, USA).

### 1.3.3 Hairpin Oligonucleotide Nomenclature

Hairpin oligonucleotides are designated by a three number designation. The first number is the total length of the oligonucleotide in nucleotides, the second number is the length of the duplex portion of the hairpin in base pairs, the third number is the length of the retained promoter consensus sequence. For some oligonucleotides an additional designation indicating the number and the type of complementary sequences in the 5’ extended template is added.

## 1.4 Results

In order to determine whether T7 RNAP is able to initiate on a truncated promoter similar to the nine nucleotide promoter of the yeast mitochondrial RNAP, we systematically deleted from the 5’ end of the 20 nucleotide (−17 to +3) double stranded region on oligonucleotides containing the promoter and determined if they would label at the 3’ end of the oligonucleotide; initiate promoter-dependent, template-independent *de novo* RNA synthesis; initiate promoter-dependent, template-dependent *de novo* transcription; or fail to incorporate label under the conditions of having the T7 RNAP and a single radiolabeled ribonucleotide triphosphate present which can base pair with the first unpaired base in the template strand (5’ extension). The starting oligonucleotide for the deletion mapping was the oligonucleotide shown in Figure 2.3C. It is able to form intra- and intermolecular base pairing to produce a recessed 3’end on an extended 5’ template. Next to the 3’ end on the unpaired template are four T nucleotides that will base pair with the radiolabeled 32P-rATP used as only nucleotide triphosphate in the experiment. As the double stranded promoter region is removed the loop length is, in some cases, increased to maintain the overall length. Table 2.1 shows the results of these experiments and Figure 2.4 shows examples of the four different results.

**Figure 2.4:**
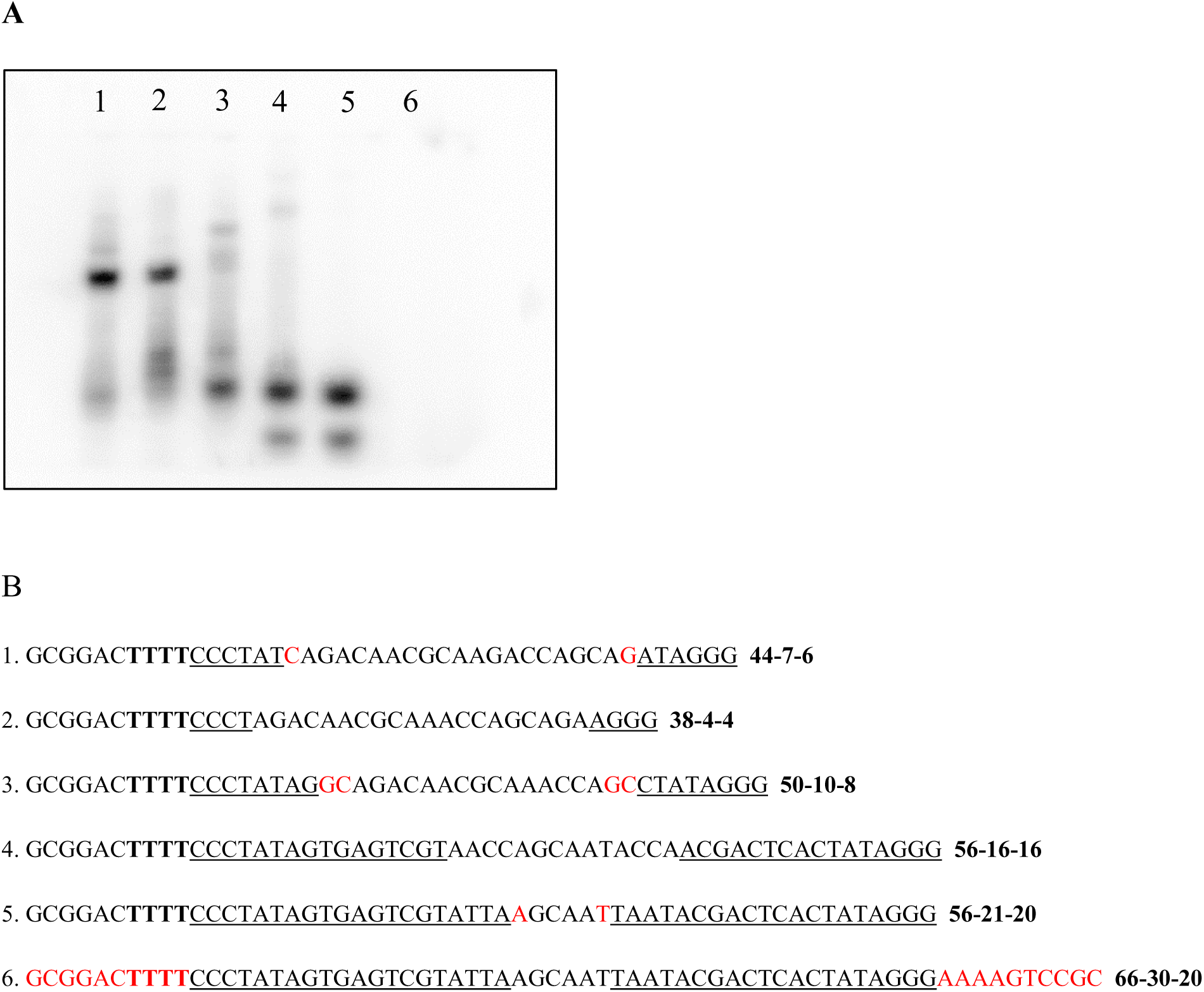
Panel A: Autoradiograph of 15% polyacrylamide gel used to separate the RNA labeled products produced from incubation of the various oligonucleotides with T7 RNA polymerase and radiolabeled rATP. The oligonucleotides used, listed from left to right (1 to 6) are 44-7-6, 38-4-4, 50-10-8, 56-16-16, 56-20-20, 66-30-20. Panel B: The DNA sequences written 5’ to 3’ for the oligonucleotides used in this experiment. Underlines indicate promoter or partial promoter sequences, red nucleotides indicate the potential for duplex beyond the promoter region, bold letters indicate potential templates for ATP radionucleotides.

**Table 2.1.**
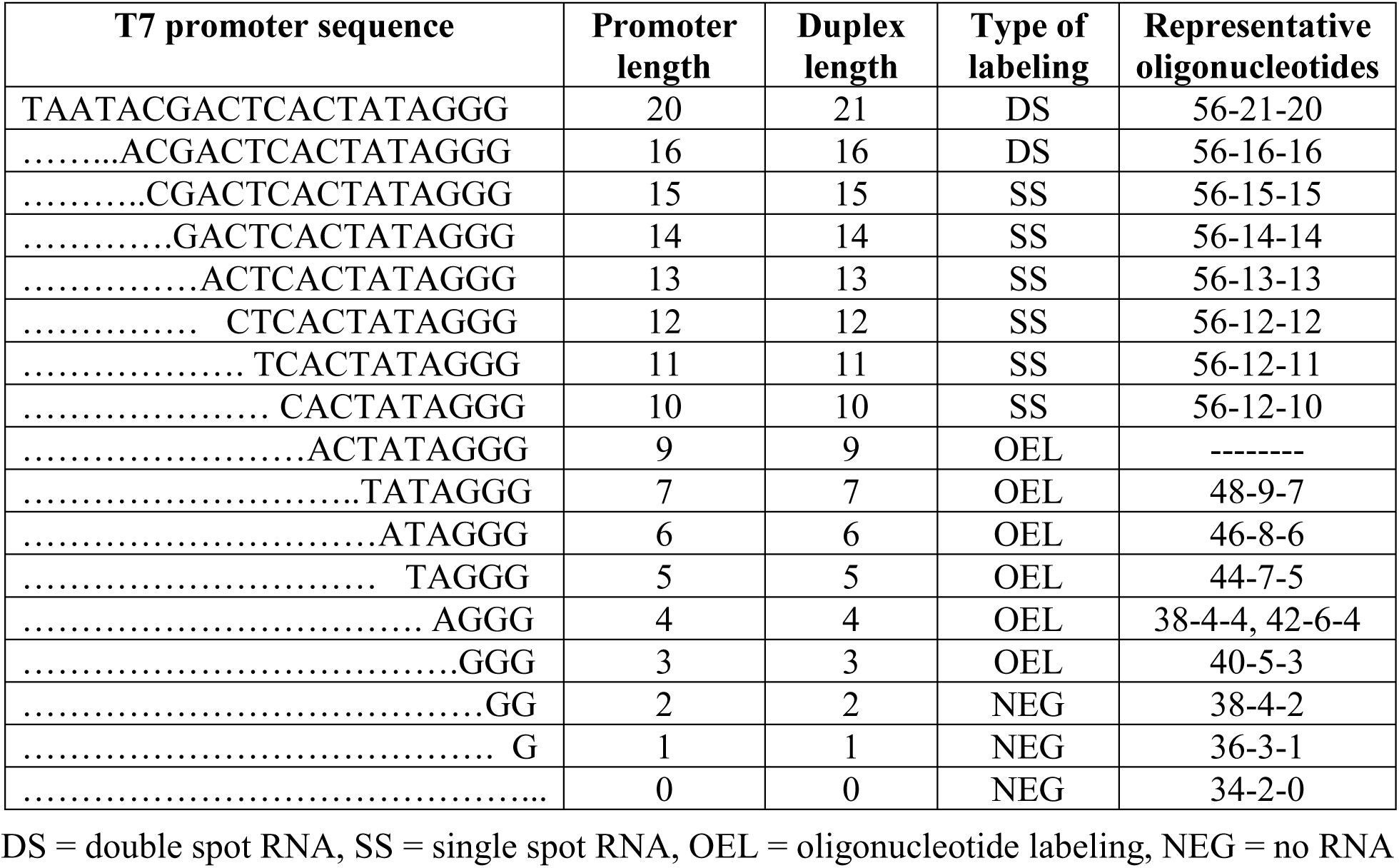
Results of the sequential deletion of the 5’ end of the double stranded promoter region of hairpin loop oligonucleotides

The oligonucleotide shown in figure 2.3C (56-21-20) has a full 20 base pair promoter sequence (− 17 to +3). With all four ribonucleotide triphosphates present T7 RNAP polymerase would be expected to initiate at +1 and make a 13 nucleotide RNA run-off transcript staring with GGG. With only rATP present T7RNAP reproducibly makes RNAs that separate upon gel electrophoresis as two high mobility, labeled spots (Figure 2.4, lane 5) designated DS for “double spot” in Table 2.1, line 1. This double spot product is RNAase sensitive (not shown) and will be called the “double spot” RNA product below.

Oligonucleotide 66-30-20 is identical to 56-21-20 except that the recessed 3’ end is extended on the template strand to produce a 30 bp double stranded region with a full promoter present and a blunt end terminus. This oligonucleotide fails to produce a “double spot” RNA product under the same conditions as with 56-21-20 (Figure 2.4, lane 6) indicating, that under the conditions used in our assay, a recessed 3’ end is required to produce the “double spot” RNA

Oligonucleotide 56-16-16 is identical to 56-21-20 except that four nucleotides are removed from the 5’ end of the double stranded promoter region to remove the AT-rich region. Eight nucleotides are added to the single stranded loop region to maintain the size of the oligonucleotide at 56 nucleotides. The −13 to +3 region of the promoter is retained (16 nucleotide promoter). Oligonucleotide 56-16-16 (Table 1, line 2) also produces a “double spot” RNA (Figure 2.4, lane 4). Oligonucleotides with 5, 6, 7, 8, 9, or 10 base pairs of double stranded promoter region deleted, to give 15 to 10 bp truncated promoters (from −12 to +3 to −7 to +3), reproducibly produce a high mobility, labeled spot designated SS (single spot) in Table 1 and fail to produce the “double spot” RNA product, indicating that a 16 bp promoter or larger is necessary for the “double spot” RNA and that the failure to produce the “double spot” RNA may allow the production of the “single spot” RNA product. These oligonucleotides lack all or part of the specificity loop region, and therefore, most of the recognition region of the promoter. Oligonucleotide 50-10-8 (−7 to +3) produces a single spot RNA product (Figure 2.4, lane 3). The “single spot” product is also sensitive to RNAase (not shown) and will be designated the “single spot” RNA product below.

Oligonucleotides with 11, 12, 13, 14, 15, 16 and 17 base pairs of 5’ double stranded promoter region deleted, to give 3 to 9 bp truncated promoters (from −6 to +3 to +1 to +3), reproducibly add a labeled nucleotide to the 3’ end of the oligonucleotide, designated OEL (oligonucleotide end labeling) in Table 2.1, and fail to produce “single spot” RNA product indicating that end labeling may prevent, or compete with, the production of “single spot” RNA. These oligonucleotides have a double stranded region with a recessed 3’ end equal to the truncated promoter length. Oligonucleotides 44-7-6 (−3 to +3) and 38-4-4 (+1 to +3) label the 3’ end of the oligonucleotide (Figure 2.4, lanes 1 and 2).

Oligonucleotides with 18, 19, and 20 base pairs of 5’ double stranded promoter region deleted, to give 0 to 2 bp truncated promoters (+2 and +3, just +3, none), do not produce an RNA product or label the 3’ end of the oligonucleotide (data not shown).

### 1.4.1 Characterization of the “double spot” RNA product

An RNA product remarkably similar to the “double spot” has been observed by Ling et al. (1989) using T7 RNAP with a full double strand promoter with four ribonucleotide triphosphates present. They ascribed this RNA product to abortive initiation. To determine if the “double spot” is abortive initiation, we examined the dependence of “double spot” on both template and promoter. In order to test the effect of the template portion of the nucleotide on “double spot” RNA production, the number of T nucleotides in the template portion of 56-21-20 were varied from 0 to 8 or substituted with A nucleotides. All of these modified oligonucleotides were able to produce the double spot RNA product (Figure 2.5), indicating that double spot RNA synthesis is template independent.

**Figure 2.5.**
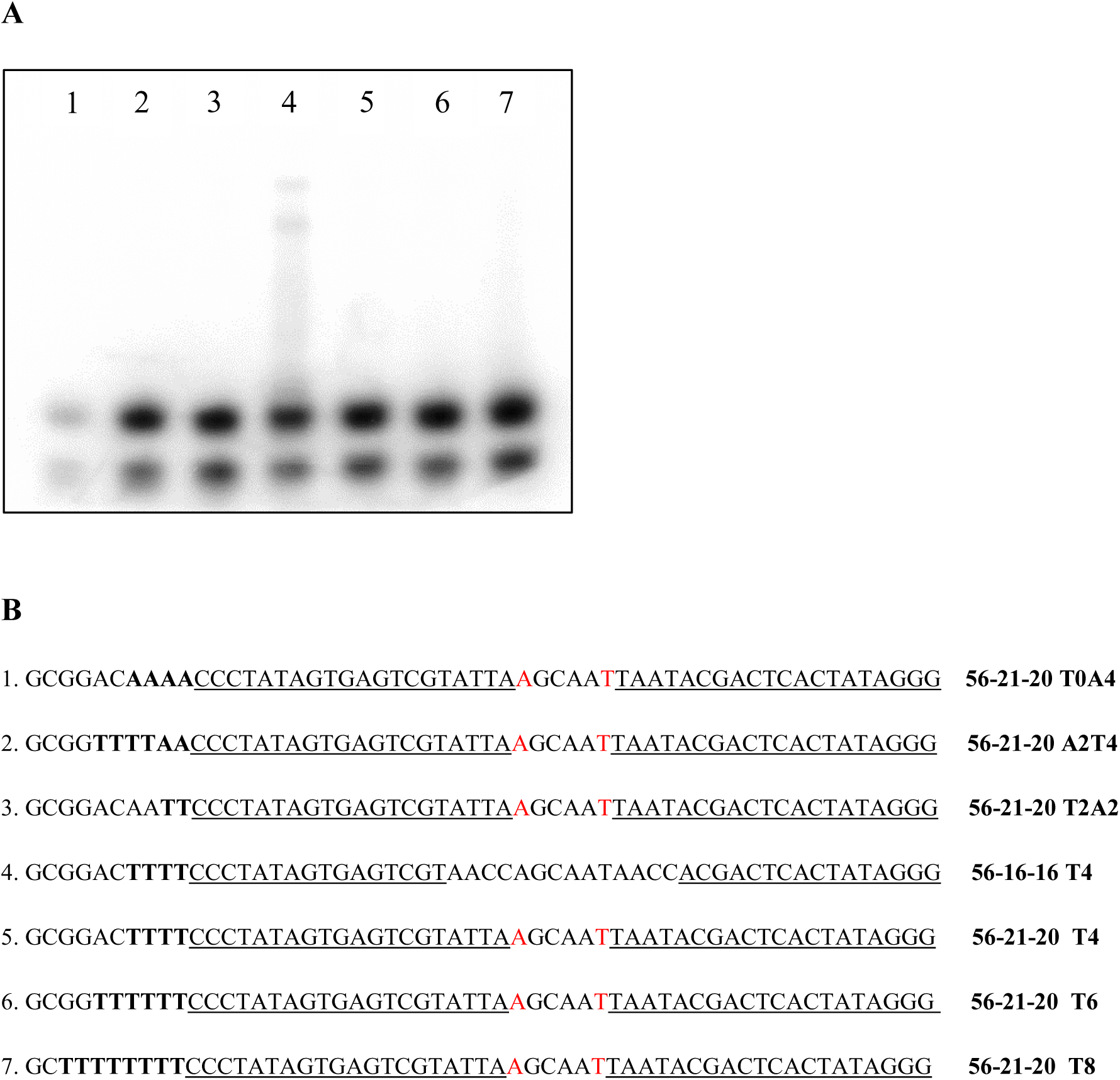
Panel A: Autoradiograph of 15% polyacrylamide gel showing double spot RNA production after incubation with T7 RNA polymerase and radiolabeled rATP of the 56-21-20 and 56-16-16 oligonucleotides modified in their template region. Panel B: The DNA sequences written 5’ to 3’ for the oligonucleotides used in this experiment. Underlines indicate double stranded promoter regions or partial double stranded promoter sequences, red nucleotides indicate the potential for duplex beyond the promoter region, bold letters indicate potential templates for ATP radionucleotides.

To determine if promoter sequences are necessary for the production of “double spot” RNA product, scrambled promoter sequences were substituted for the promoter sequence. While the classic “double spot” nucleotides 56-21-20, and 56-14-14 produced “double spot” RNA product, the oligonucleotides with scrambled promoter sequences did not produce “double spot” RNA product (Figure 2.6), indicating that double spot production is dependent on promoter sequence

**Figure 2.6.**
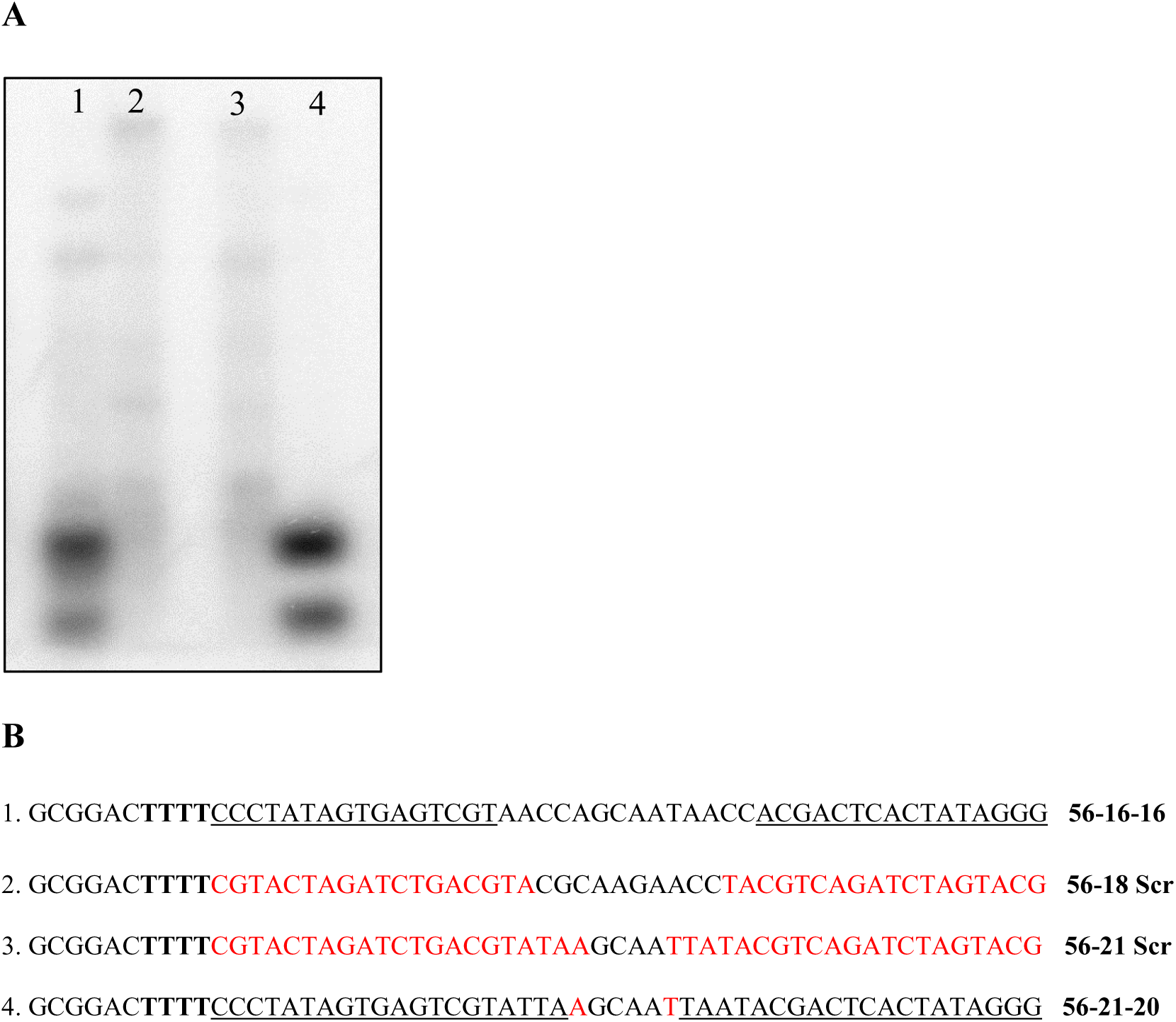
Panel A: Autoradiograph of 15% polyacrylamide gel showing double spot RNA production after incubation with T7 RNA polymerase and radiolabeled rATP with the 56-21-20 and 56-16-16 oligonucleotides modified to have scrambled promoter sequences. Panel B: The DNA sequences written 5’ to 3’ for the oligonucleotides used in this experiment. Underlines indicate double stranded promoter regions or partial double stranded promoter sequences, red nucleotides indicate the potential for duplex beyond the promoter region, bold letters indicate potential templates for ATP radionucleotides.

These results indicating that “double spot” RNA production is template independent and promoter dependent are consistent with the report that “double spot” RNA is abortive initiation. 5’ end deletions of 4, 5, and 6 nucleotides (oligonucleotides 56-16-16, 56-15-15, and 56-14-14) can produce “double spot” RNA (Figure 2.7). Oligonucleotide 56-15 has the AT-rich recognition sequence 5’-TAATA-3’ substituted with 5’-TATAT-3’, a similar sequence. This substitution has little effect on “double spot” RNA production (Figure 2.7, lane 3). To determine if a GC–rich sequence can be tolerated as a substitute for the AT-rich recognition sequence, the sequence CGCGG was substituted in 56-15-5GC. This oligonucleotide produces only a very small amount of double spot” DNA (Figure 2.7, lane 4), indicating that while the primary sequence of the AT-rich region is not critical, the base composition of the region should be AT-rich to give optimum double spot RNA production.

**Figure 2.7.**
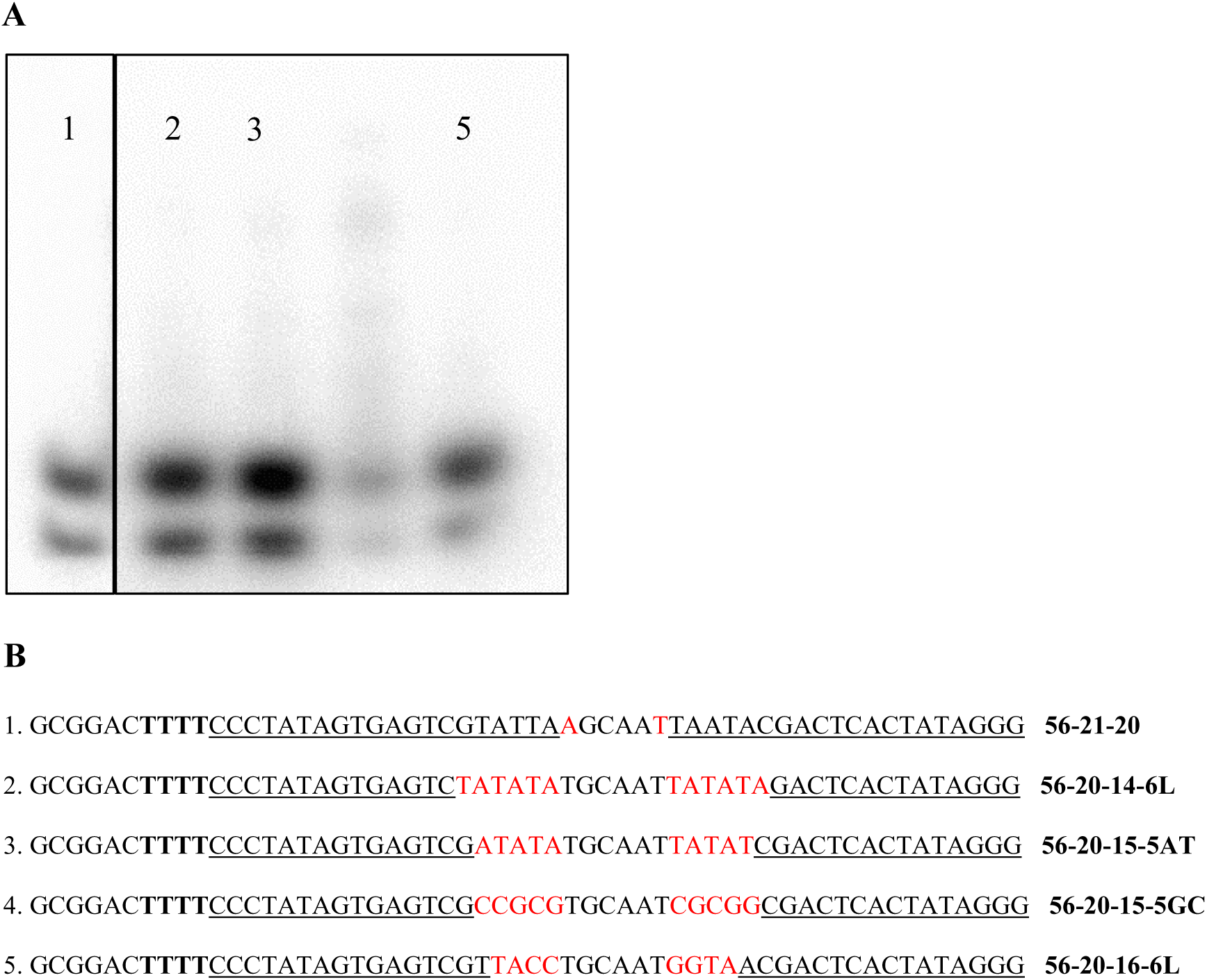
Panel A: Autoradiograph of 15% polyacrylamide gel showing double spot RNA production after incubation with T7 RNA polymerase and radiolabeled rATP with the 56-21-20, 56-20-16, 56-20-15 oligonucleotides. The 56-20-16 and 56-20-15 oligonucleotides have substituted bases in the AT-rich binding region. Panel B: The DNA sequences written 5’ to 3’ for the oligonucleotides used in this experiment. Underlines indicate double stranded promoter regions or partial double stranded promoter sequences, red nucleotides indicate the potential for duplex beyond the promoter region, bold letters indicate potential templates for ATP radionucleotides.

Our interpretation of these results is that “double spot” RNA production is equivalent to abortive *de novo* initiation at the recessed 3’ end. A complete 20 nucleotide promoter (56-21-20; −17 to +13) or nearly complete 5’ truncated promoters (56-16-16, 56-20-16, 56-20-15, and 56-20-14; - 13, −12, −11 to +13) bind strongly enough so that it cannot leave the promoter during promoter clearance during the transition from initiation to elongation. The presence of only a single ribonucleotide triphosphates also prevents elongation. The transcripts produced prior to promoter clearance produce the abortive initiation products that we designate “double spot” RNA

### 1.4.2 Oligonucleotide 3’ end labeling

The labeling of oligonucleotides with recessed 3’ ends by T7 RNA polymerase has been reported and characterized by Sarcar & Miller (manuscript submitted) as a type of DNA editing. They report a promoter-independent, template dependent, recessed 3’ end dependent addition of rNTPs to the 3’ end of the DNA. A ribonucleotide is added to the 3’ end when the 3’ end is positioned next to a single stranded region where the first unpaired nucleotide is complementary to the labeling ribonucleotide triphosphate. While this process absolutely requires a double stranded region with a recessed 3’ end, it is limited by the length of the base paired region.

Oligonucleotides with duplex regions shorter that 3 base pair or longer than about 8 base pairs, depending on their base composition, do not label. In our initial detection of oligonucleotide 3’ end labeling, we observed labeling of oligonucleotide with duplex potentials of 3 - 9 base pairs (Table 2.1). However, in this initial screen the partial promoter length was equal to, or approximately equal, to the duplex length of the hairpin oligonucleotide (Table 2.1). To uncouple these two parameters, we made oligonucleotides with a constant 20 bp duplex length, but with variable promoter lengths (see for example, 56-20-16, 56-20-15, 56-20-14, Figure 2.7).

These showed “double spot” or “single spot” RNA production depending on partial promoter length, but did not show end labeling since the duplex length exceeded the 9 nucleotide length necessary for end labeling. Conversely, oligonucleotides without promoter sequences, but hairpin oligonucleotides with duplex lengths between 3 and 9 base pairs and recessed 3’ends with complementary sequences, labeled efficiently.

Figure 2.8 shows oligonucleotides with constant loop length (15 nucleotides) and template length (10 nucleotides) and increasing duplex length from 2 bp to 10 bp. Oligonucleotides with duplexes 4 to 8 bp label efficiently; oligonucleotides with duplexes of 3 and 9 bp label slightly. Oligonucleotides with smaller or larger duplex length (2 bp and 10 bp) did not 3’ end label (Figure 2.8).

**Figure 2.8.**
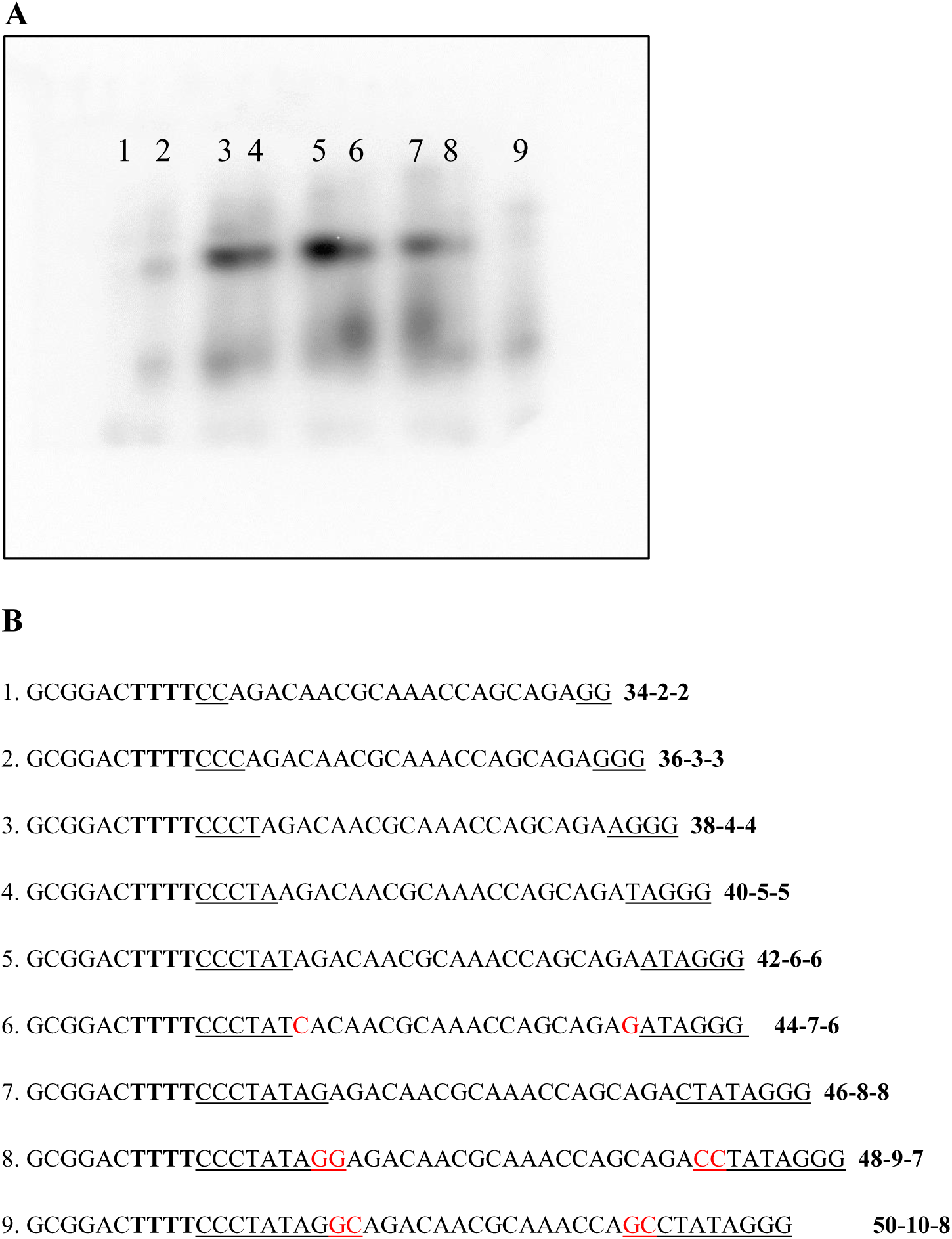
Panel A: Autoradiograph of 15% polyacrylamide gel showing oligonucleotide 3’ end labeling after incubation with T7 RNA polymerase and radiolabeled rATP. Each hairpin oligonucleotide has a duplex length one bp longer than the previous oligonucleotide. Panel B: The DNA sequences written 5’ to 3’ for the oligonucleotides used in this experiment. Underlines indicate double stranded promoter regions or partial double stranded promoter sequences, red nucleotides indicate the potential for duplex beyond the promoter region, bold letters indicate potential templates for ATP radionucleotides.

To determine template dependence in oligonucleotide 3’ end labeling, the 5’ extension/template of the 44-7-6 oligonucleotide was systematically altered. Substitution of A nucleotides for T nucleotides in the labeling site resulted in no labeling (Figure 2.9, lanes 1 and 7), indicating that the labeling site next to the 3’ end must base pair with the radionucleotide triphosphate (rATP) used in the assay. Lanes 2, 3, 4, 5, and 6 of Figure 2.9 have oligonucleotides with a variable number of T nucleotides in the labeling site giving the potential for addition of multiple A nucleotides to the 3’ end of the oligonucleotide. Although at least one A nucleotide is added in each case since labeling occurs, it is unlikely that multiple A nucleotides are added. This conclusion is based on the fact that the intensity of labeling is essentially equal, except for lane 3 where labeling intensity is decreased not increased for an eight T nucleotide template, and the migration of the labeled oligonucleotide is not changed.

**Figure 2.9.**
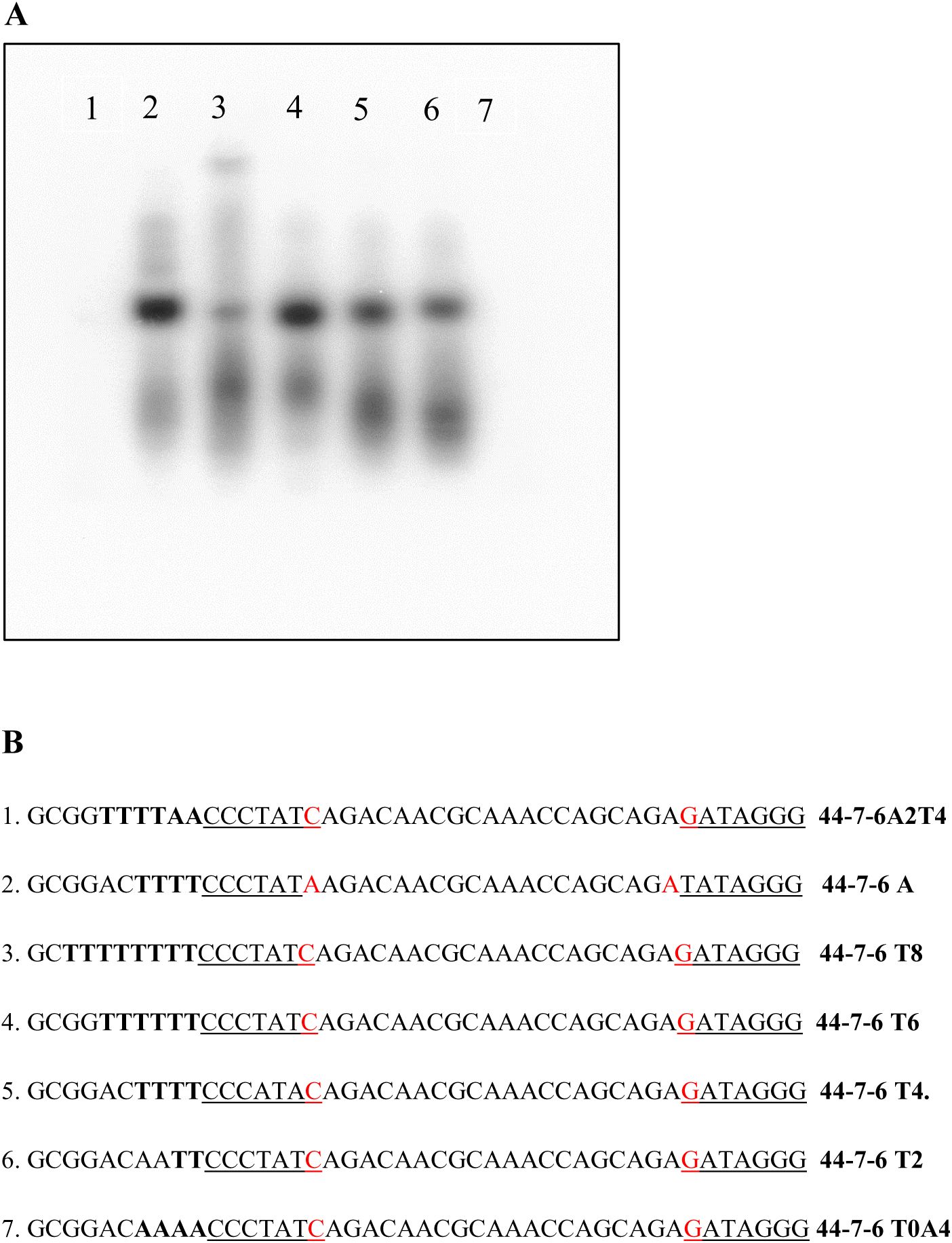
Panel A: Autoradiograph of 15% polyacrylamide gel showing oligonucleotide 3’ end labeling after incubation with T7 RNA polymerase and radiolabeled rATP. Each hairpin oligonucleotide has a constant length (44 nucleotides), constant duplex length (7 bp), and constant loop length (15 nucleotides). The template 5’ extension is a constant length (10 nucleotides) but has a variable number of T nucleotides at the labeling site or A nucleotide substitutions at the site. Panel B: The DNA sequences written 5’ to 3’ for the oligonucleotides used in this experiment. Underlines indicate double stranded promoter regions or partial double stranded promoter sequences, red nucleotides indicate the potential for duplex beyond the promoter region, bold letters indicate potential templates for ATP radionucleotides.

Addition of multiple nucleotides to the 3’ end of the oligonucleotide would be expected to change the migration of the oligonucleotide in gel electrophoresis. Interestingly, there seems to be concurrent end labeling and *de novo* initiation in the autoradiograph of Figure 2.9. The *de novo* initiation does seem to elongate up to 8 nucleotides (see the similar experiment in Figure 2.13 below).

### 1.4.3 Charaterization of the “single spot” RNA product

Under the conditions used in these experiments, it was not expected that the oligonucleotides would be capable of *de novo* intiaition, since promotor dependent *de novo* intiation requires three G nucleotides for intiation and rGTP is not provided in these experiments. Nonetheless, a small RNA, presumably 5’-pApApApA-3’, was produced when four T nucleotides are positioned next to the recessed 3’ end in hairpin oligonucleotides with 5’ truncated promoter sequences between 10 and 15 nucleotides.

In Figure 2.10 six oligonucleotides with promoter lengths between 10 and 15 are able to produce “single spot” RNAs, but scrambling of promoter sequences completely eliminates “single spot” RNA production (Figure 2.11), indicating that this activity is dependent on a partial promoter sequence. While the duplex region of the hairpin oligonucleotides requires a partial promoter sequence, variable loop lengths do not affect the single spot production (Figure 2.12). However, the single spot production is dependent on the 5’ extension/template region of the oligonucleotide (Figure 2.13). Oligonucleotide 56-12-10 A4 (Figure 2.13, lane 1) has A substitutions for the template T nucleotides and does not produce RNA. This indicates “single spot” RNA production is template dependent and requires a complementary base in the site next to the 3’ end.

**Figure 2.10.**
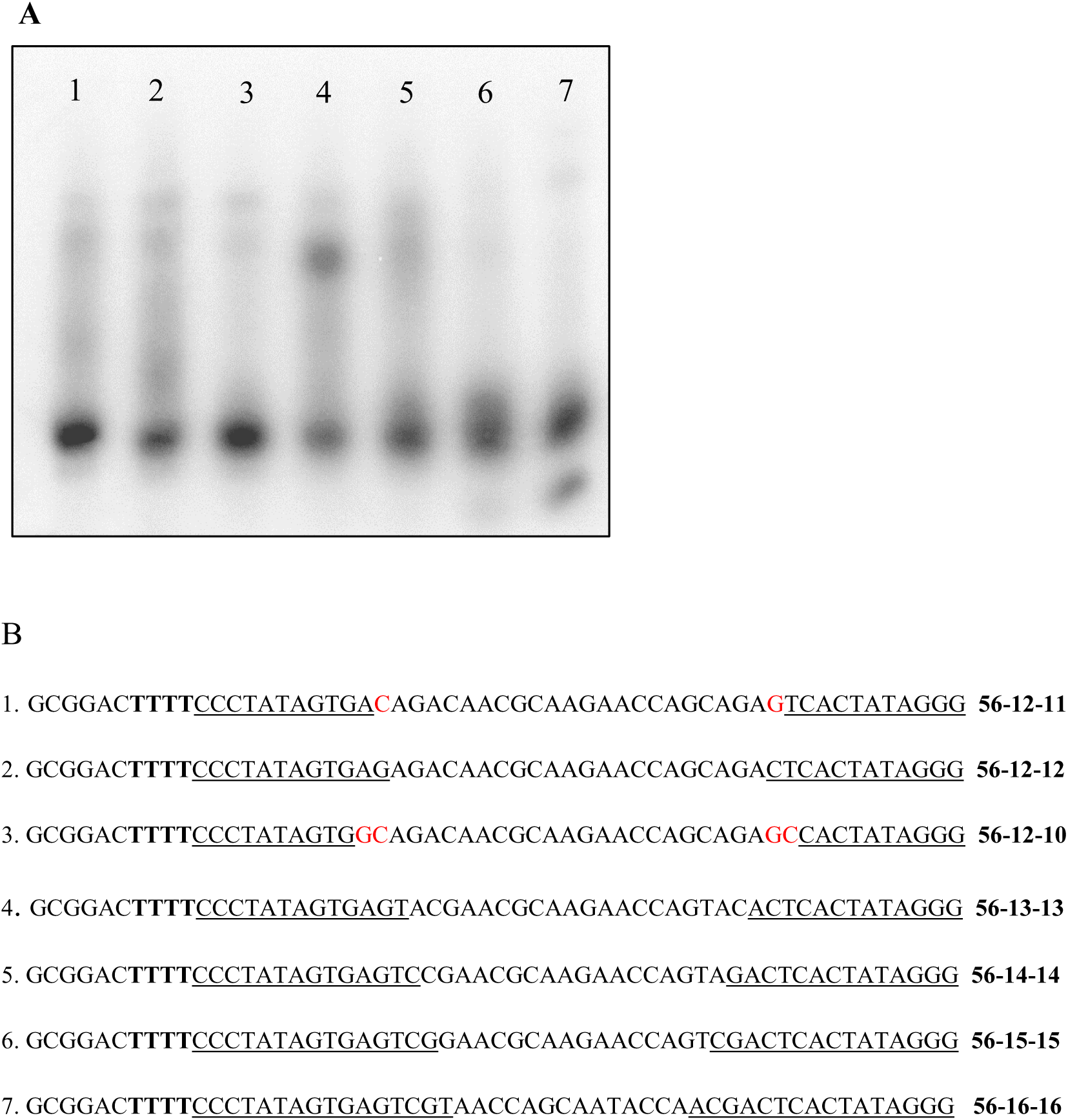
Panel A: Autoradiograph of 15% polyacrylamide gel showing oligonucleotide “single spot” RNA production with variable length partial promoter sequences (10 – 15 bp) labeling after incubation with T7 RNA polymerase and radiolabeled rATP. Oligonucleotides with partial promoter sequences longer than 15 bp (56-16-16) produce “double spot” RNA. Panel B: The DNA sequences written 5’ to 3’ for the oligonucleotides used in this experiment. Underlines indicate double stranded promoter regions or partial double stranded promoter sequences, red nucleotides indicate the potential for duplex beyond the promoter region, bold letters indicate potential templates for ATP radionucleotides.

**Figure 2.11.**
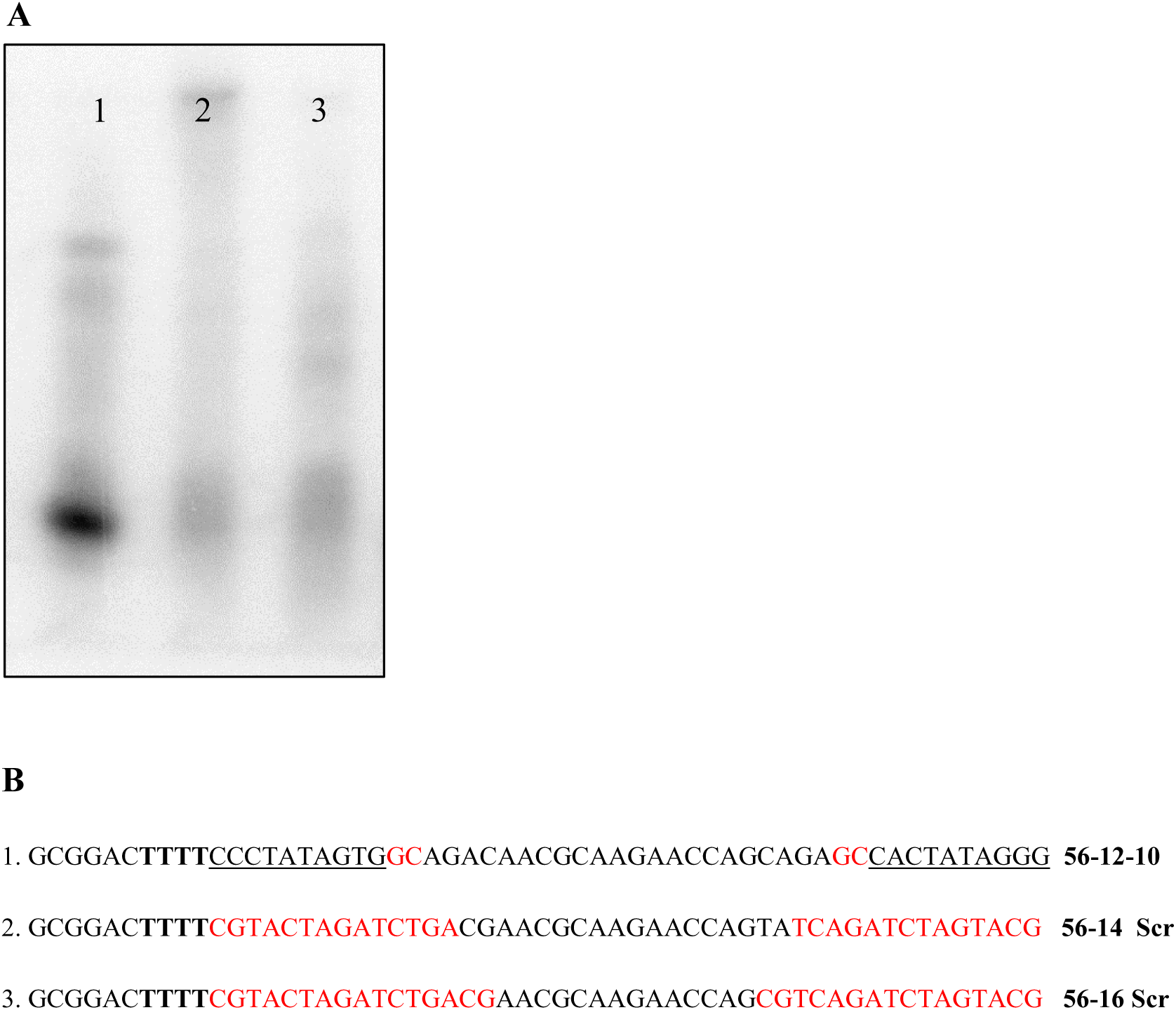
Panel A: Autoradiograph of 15% polyacrylamide gel showing single spot RNA production after incubation with T7 RNA polymerase and radiolabeled rATP. Oligonucleotide 56- 12-10 (lane 1) has a 10 bp partial promoter sequence and produces “single spot” RNA. Oligonucleotides 56-14 Scr and 56-16 Scr contain scrambled promoter sequences and do not produce single spot RNA. Panel B: The DNA sequences written 5’ to 3’ for the oligonucleotides used in this experiment. Underlines indicate double stranded promoter regions or partial double stranded promoter sequences, red nucleotides indicate the potential for duplex beyond the promoter region, bold letters indicate potential templates for ATP radionucleotides.

**Figure 2.12.**
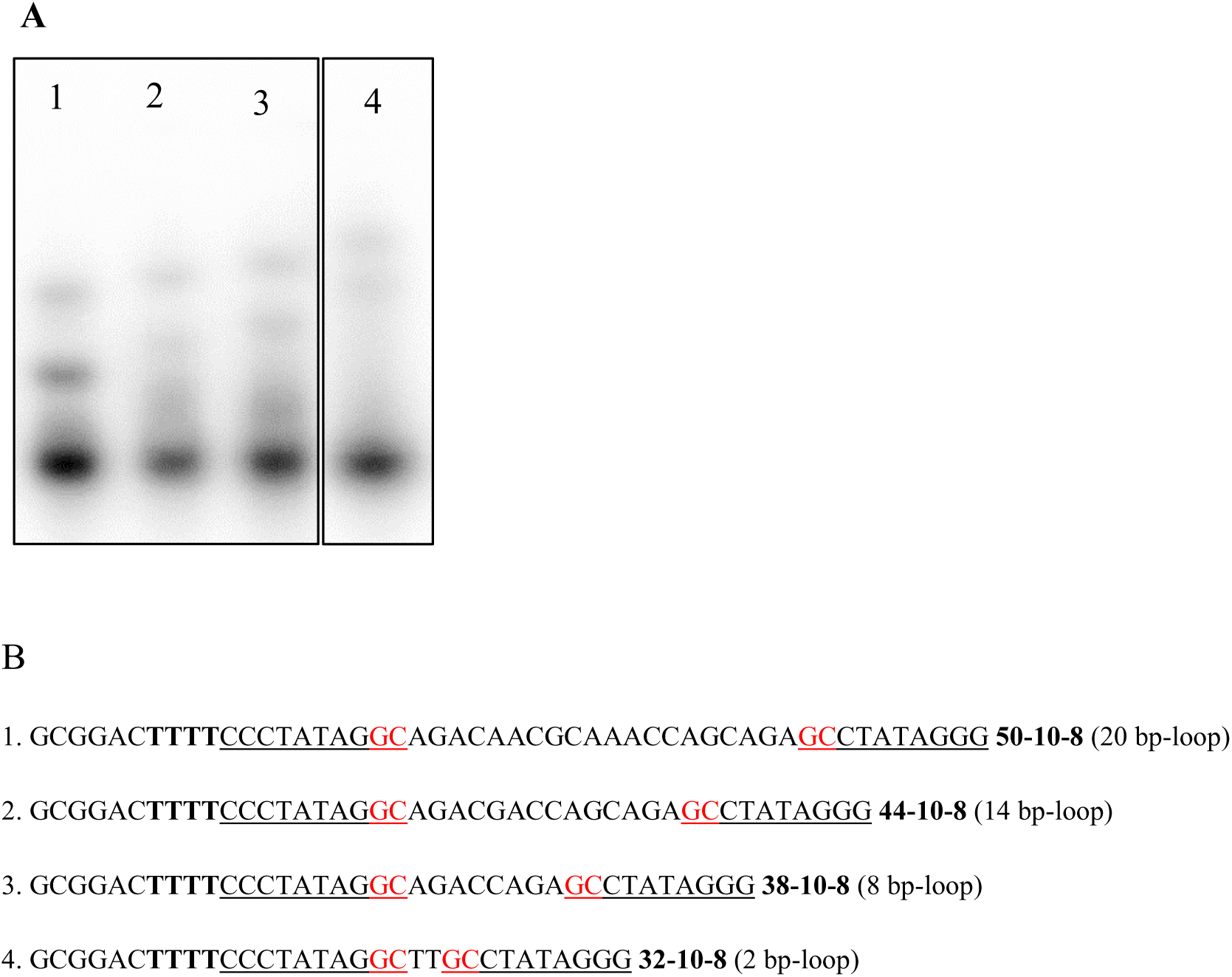
Panel A: Autoradiograph of 15% polyacrylamide gel showing “single spot” RNA after incubation with T7 RNA polymerase and radiolabeled rATP. Each hairpin oligonucleotide has a duplex length of 10 bp, a 5’ truncated promoter to 8 bp, and a 10 nucleotide 5’extension. Each hairpin oligonucleotide has a different loop length. Panel B: The DNA sequences written 5’ to 3’ for the oligonucleotides used in this experiment. Underlines indicate double stranded promoter regions or partial double stranded promoter sequences, red nucleotides indicate the potential for duplex beyond the promoter region, bold letters indicate potential templates for ATP radionucleotides.

The remaining oligonucleotides in Figure 2.13 have 2, 4, 5, 6, and 8 T nucleotides in the template region next to the 3’ end. The RNAs made from these oligonucleotides correspond in length to the template, indicating that they can initiate template-dependent *de novo* RNA synthesis from a recessed 3’ end when a partial promoter is present and elongate the RNA on the template. This *de novo* initiation site is located at +4 in contrast to the classical site of transcription initation at +1

**Figure 2.13.**
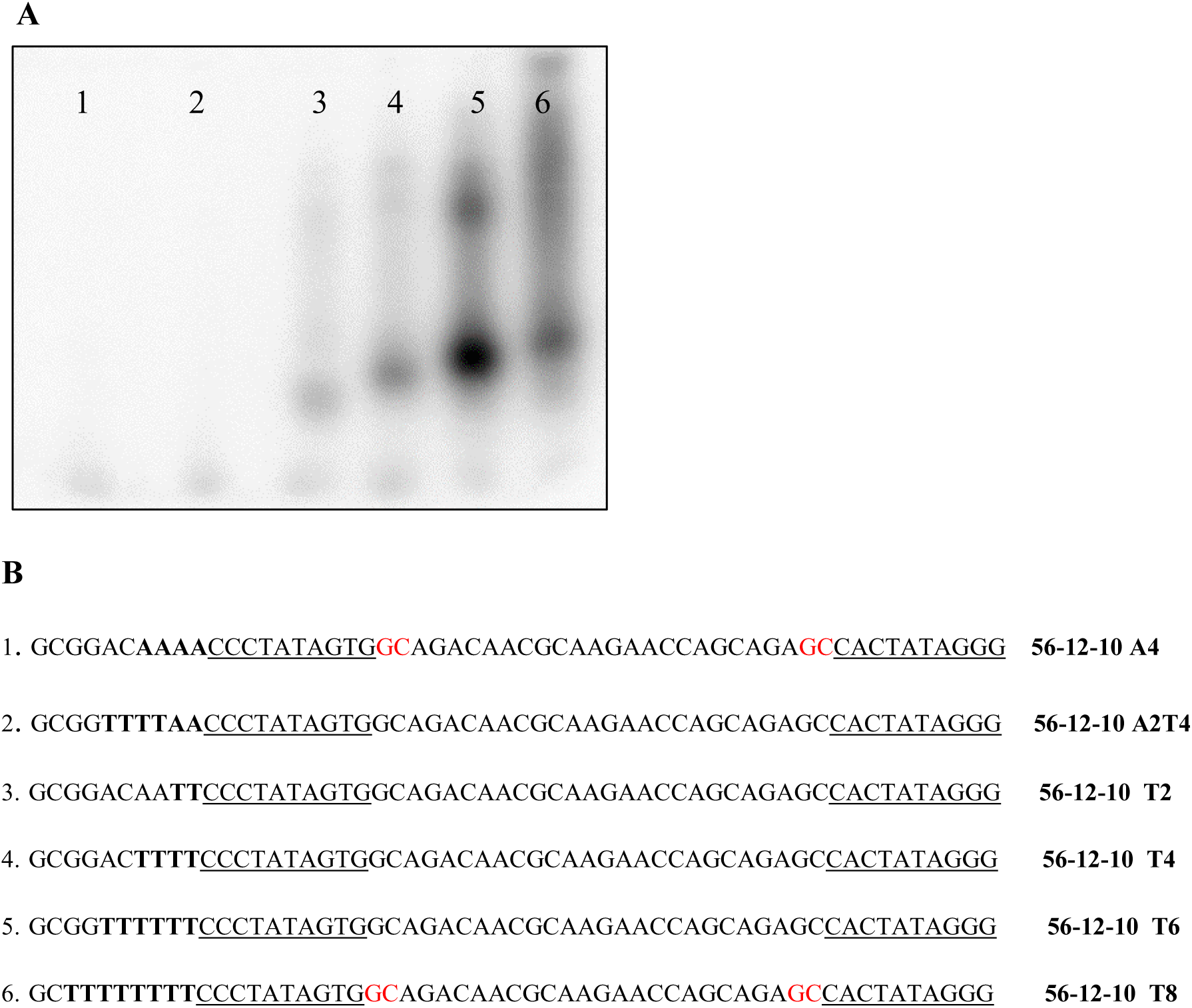
Panel A: Autoradiograph of 15% polyacrylamide gel showing *de novo* initiation with template directed elongation from oligonucleotides with recessed 3’ ends and partial promoter sequences after incubation with T7 RNA polymerase and radiolabeled rATP. Panel B: The DNA sequences written 5’ to 3’ for the oligonucleotides used in this experiment. Underlines indicate double stranded promoter regions or partial double stranded promoter sequences, red nucleotides indicate the potential for duplex beyond the promoter region, bold letters indicate potential templates for ATP radionucleotides.

## 1.5 Discussion

When presented with an oligonucleotide which can form a hairpin loop with a recessed 3’ end and a single ribonucleotide triphosphate, T7RNAP displays several novel activities depending on the amount of the promoter sequence retained in the duplex region of the hairpin. When a full promoter (20 nucleotides, −17 to +3) or up to a four base pair 5’ deletion of the full double stranded promoter (16 nucleotide, −13 to +3) is used in these circumstances a double spot, abortive transcript is produced. Although this *de novo* initiation can occur if a recessed 3’ end is present, it is template independent and produces the same double spot pattern independent of the sequence of the 5’ extension. This is presumably the result of strong promoter binding due to the retention of the −12 to −5 region of the promoter which is bound by the specificity loop.

Five to ten base pair deletions of the 5’ end of the double stranded promoter result in hairpin oligonucleotides with 15 bp (−12 to +3) to 10 bp (−7 to +3) partial promoters that can initiate *de novo* template-dependent RNA synthesis. These RNAs initiate at the first unpaired base after the recessed 3’ end at a +4 relative to classical full promoter driven transcription. These partial promoters resemble mitochondrial promoters in length, 9 nucleotides (−8 to +1), however, mitochondrial promoters initiate at +1. The minimal T7 RNAP promoter sequence able to support *de novo* initiation at a recessed 3’ end *in vivo* is 5’-CACTATAGGG-3’. Deletion of promoters beyond 10 base pairs results in the loss of *de novo* initiation. However, hairpin oligonucleotides with recessed 3’ ends and a duplex shorter than about 9 base pairs of AT–rich sequence or longer than about 3 base pairs of GC-rich sequence can add a Watson-Crick specified ribonucleotide to the 3’ end of DNA oligonucleotides (Sarcar & Miller, manuscript submitted), producing a DNA-RNA phosphodiester bond. The addition of a specific ribonucleotide to a DNA oligonucleotide constitutes an RNA editing of DNA. This activity may be similar to the co-transcriptional, non-DNA templated, insertional RNA editing activity observed for the mtRNAP in mitochondria of myxomycetes (D. Miller, Padmanabhan, & Sarcar, 2017; M. L. Miller & Miller, 2008; Visomirski-Robic & Gott, 1997). The evolutionary relationship of the single subunit polymerases is obscure even though they are clearly related through highly conserved motifs. DNA-dependent DNA polymerases and RNA-dependent DNA polymerases utilize RNA or DNA primers to initiate DNA synthesis at specific sites determined by the complementarity of the primer. Although primer extension is also used during elongation by RNA polymerases, they have uniquely evolved the ability to initiate *de novo* transcription through promoter recognition. Here we show that in the correct sequence context (partial promoters), primers can be used to specify *de novo* initiation of transcription *in vitro.* This activity may represent a transitional activity from primer-initiated nucleic acid synthesis to promoter-directed initiation in the evolution of RNA polymerases.

In any case, the discovery of primer-specified *de novo* initiation *in vitro* suggests a method of synthesizing from oligonucleotide templates, RNAs without G nucleotides at the 5’ end and without non-DNA templated nucleotides at the 3’ end. Experiments to determine whether the use of partial promoter sequences similar to mitochondrial promoters will work with T7 polymerase *in vivo* are in progress

